# Evolutionary changes of an intestinal *Lactobacillus reuteri* during probiotic manufacture

**DOI:** 10.1101/567974

**Authors:** Lázaro López, Paúl Cardenas, María B. Prado, Gabriel Trueba

## Abstract

Probiotic bacteria are frequently used to treat intestinal (and other types of diseases) or to improve health, however little is known about the evolutionary changes of these bacteria during probiotic manufacture. It has been observed that when bacteria adapt to a new environment, they lose the skills to thrive in the original niche. In this study, a strain of *Lactobacillus reuteri* was isolated from the duodenum of a mouse and subjected to 150 serial passes in milk to simulate industrial propagation of probiotic bacteria. The strain adapted to milk outperformed its ancestor in milk, but it showed reduced aptitude to grow culture media, and possibly in mouse intestines. Bacterial adaptation to milk seemed to select a number of non-synonymous mutations in metabolic genes.

**IMPORTANCE:** Bacteria in the microbiota must endure hostile conditions including antagonism from other members of the microbial community. Little is known about the evolution of probiotic bacteria in the dairy industry. We show that repeated growth of *L. reuteri* in milk results in reduced ability to grow in culture medium and possibly colonize the mammalian intestine. Our results contribute to the notion that probiotic bacteria colonize only transiently intestines.

## INTRODUCTION

The International Scientific Association for Probiotics and Prebiotics describes probiotics as “live microorganisms that, when administered in adequate amounts, confer a benefit to the health of the host” (1). There are some properties that are desirable in a probiotic bacteria: inherent benefit to the host, lack of antibiotic resistance genes and survival in the intestine (2). Microorganism entering the microbiota must cope with bacterial toxins and antimicrobial systems produced by resident bacteria in the intestine (3–6).

The genus *Lactobacillus* encompasses different species (*L. rhamnosus, L. acidophilus, L. casei, L. bulgaricus*, and *L. reuteri*) which are normally found in the human intestine and often used in probiotics (7, 8). *L. reuteri* was isolated for the first time in 1962 and it has been described as a heterofermentative anaerobic bacterium which tolerates a wide range of pH. *L. reuteri* has coevolved with different animal hosts (including humans) and diversified into distinct phylogenetic lineages (46). In the microbiota, *L. reuteri* inhibits other bacteria and even secretes antimicrobial compounds (8, 9).

It is estimated that in the USA around 3.9 million people consume prebiotic/ probiotic supplements, probiotics are one of the most consumed dietary supplements (10) and important therapeutic products (8, 11–15). Nevertheless, there are many questions about the efficacy of these products in the medical field (16). One of the main questions is how long the probiotic bacteria can linger in the intestine.

The survival of a microorganism in the digestive tract requires tolerance to low pH (stomach), antimicrobial peptides, biliary salts, immune response, etc. One of the most important barriers that entering bacteria have to overcome is the antagonism and competition from the native members of the intestinal microbiota, some of which produce antibiotic compounds, antimicrobial peptides, secretion systems, etc. (6, 17, 18).

When bacteria adapt to a new environment, they lose the skills necessary to thrive in the original niche (19). Numerous evolutionary studies have been designed to understand how populations adapt to new specific environmental conditions, like nutrients, temperature, parasites, competition and other environmental stressors (20–24). It is assumed that most adaptations to the new condition are associated with the loss of adaptation to the original (20, 24). Adaptation occurs through the selection of mutants that deal better with different substrates, temperatures, pH values, etc. (19, 25) We studied the evolutionary changes observed in an *L. reuteri* isolate after 150 passages milk.

## RESULTS

### Competitions between Milk adapted and Ancestral strains in Milk and MRS medium

In order to identify the *L. reuteri* strain from other lactobacilli in the intestine, we selected a rifampin-resistant mutant (*L. reuteri*^RR^). *L. reuteri*^RR^ strain after 150 passes in sterile cow milk outperformed (66%) the *L. reuteri*^RR^ ancestor (relative fitness = 1.66) and the original ancestor (rifampicin sensitive) in 33% (relative fitness = 1.33) in milk. While the *L. reuteri*^RR^ strain without passes in sterile cow milk had a fitness relative to the original ancestor (rifampicin sensitive) of 0.80 in milk (figure 1). On the other hand, the growth capacity tests in MRS medium showed that *L. reuteri*^RR^ strain without milk passages was 5% less efficient than the rifampicin sensitive ancestor (fitness relative to ancestor = 0.95), whereas *L. reuteri*^RR^ strain adapted to the milk grew 16% less than its *L. reuteri*^RR^ ancestor in MRS (fitness relative to the ancestor = 0.84) (table 1).

**Figure 1.**
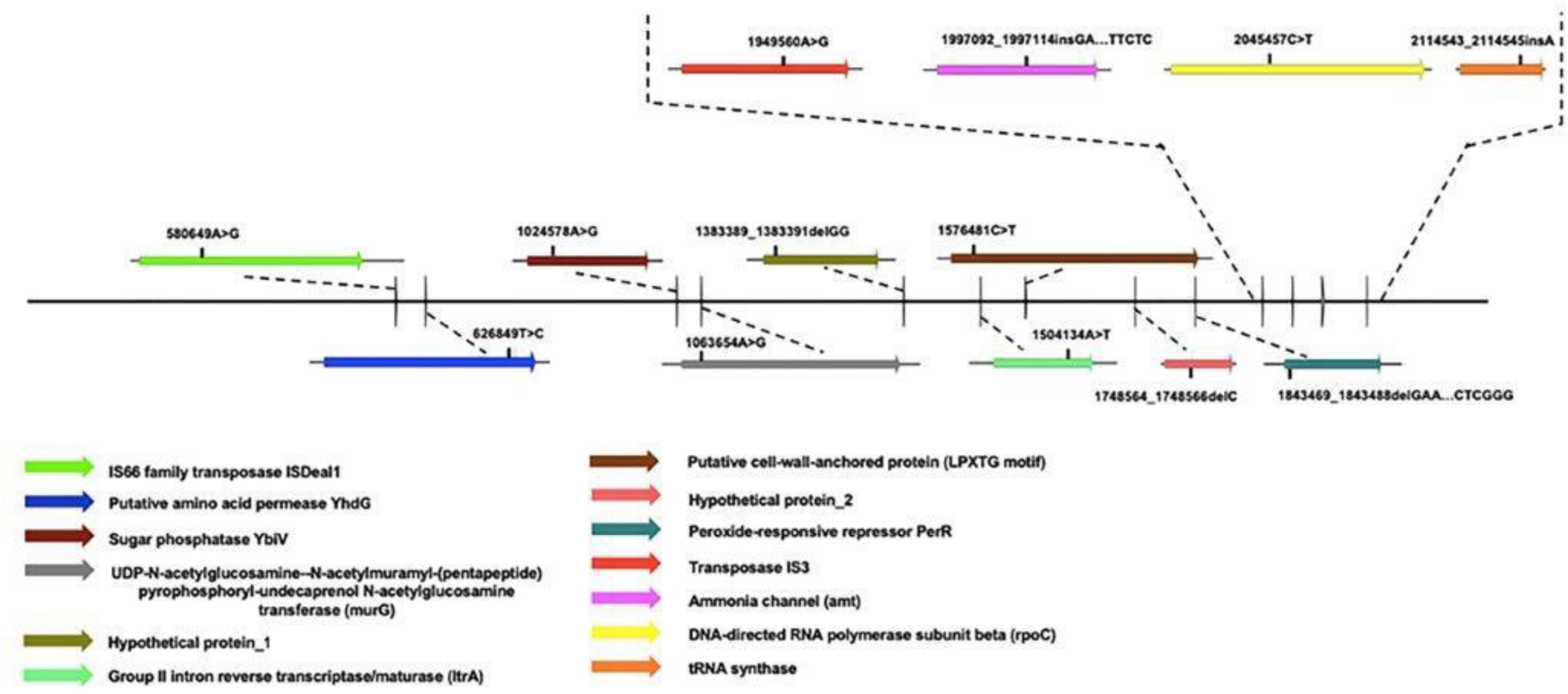
Competition in milk between without previous passes in milk (MRR (0)) and *Lactobacillus reuteri* rifampicin resistant mutant after 150 passes in milk (MRR (150)), against the original *Lactobacillus reuteri* rifampicin sensitive ancestor (red line). * Fitness relative to the ancestor expressed as a percentage

**Table 1.** Competitions between mutant strains resistant to rifampicin and their ancestor, in MRS medium.

| Strain | Relative Fitness | Percentage |
| --- | --- | --- |
| MRR(0)* | 0.95 | -5 % |
| MRR(150)** | 0.84 | -16% |

\* MRR (0) = *Lactobacillus reuteri* mutant resistant to rifampicin without passes in milk.
\*\*MRR (150) = *Lactobacillus reuteri* mutant resistant to rifampin with 150 passes in milk.

### Intestinal colonization

A culture of *L. reuteri*^RR^ from murine intestines was successful only in 2 mice (Figure supp1), however, in a mouse receiving the *L. reuteri*^RR^ ancestral strain had 35 times higher *L. reuteri*^RR^ colony counts than the animals receiving the *L. reuteri*^RR^ strain passaged in milk (993 CFU/ml vs 28 CFU/ml).

### Whole Genome Sequencing

The alignment made in Mauve (Figure supp.1) showed that all the contigs were correctly aligned among the three analyzed genomes. Comprehensive Genome Analysis in Patrick showed that the three genomes have 2,320 Mb, 2,303 Mb and 2,296 Mb respectively distributed in 310, 264 and 315 contigs (supplementary table 3). Genes annotation allowed us to compare 1,618 proteins with functional assignments and 647 hypothetical proteins from *L. reuteri*^RR^ milk-adapted strain with the *L. reuteri*^RR^ strain. Of the coding-sequence genes analyzed 13 (0,6%) had mutations (substitutions, insertions or deletions) compared to the progenitor (table 2). Of the 13 mutations found, 8 mutations were non-synonymous. Three out of 396 metabolic genes showed non-synonymous mutations.

**Table 2.**
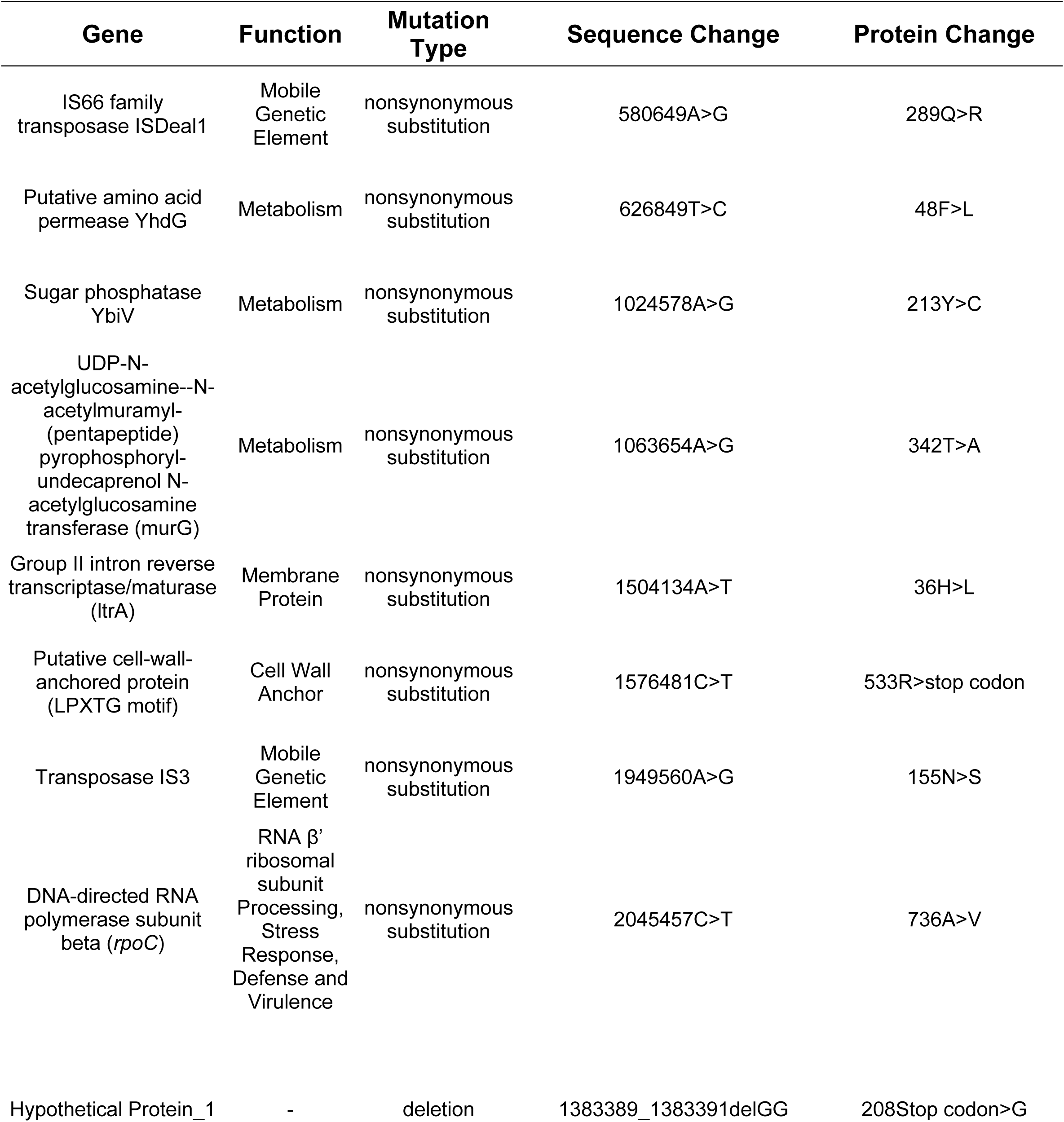
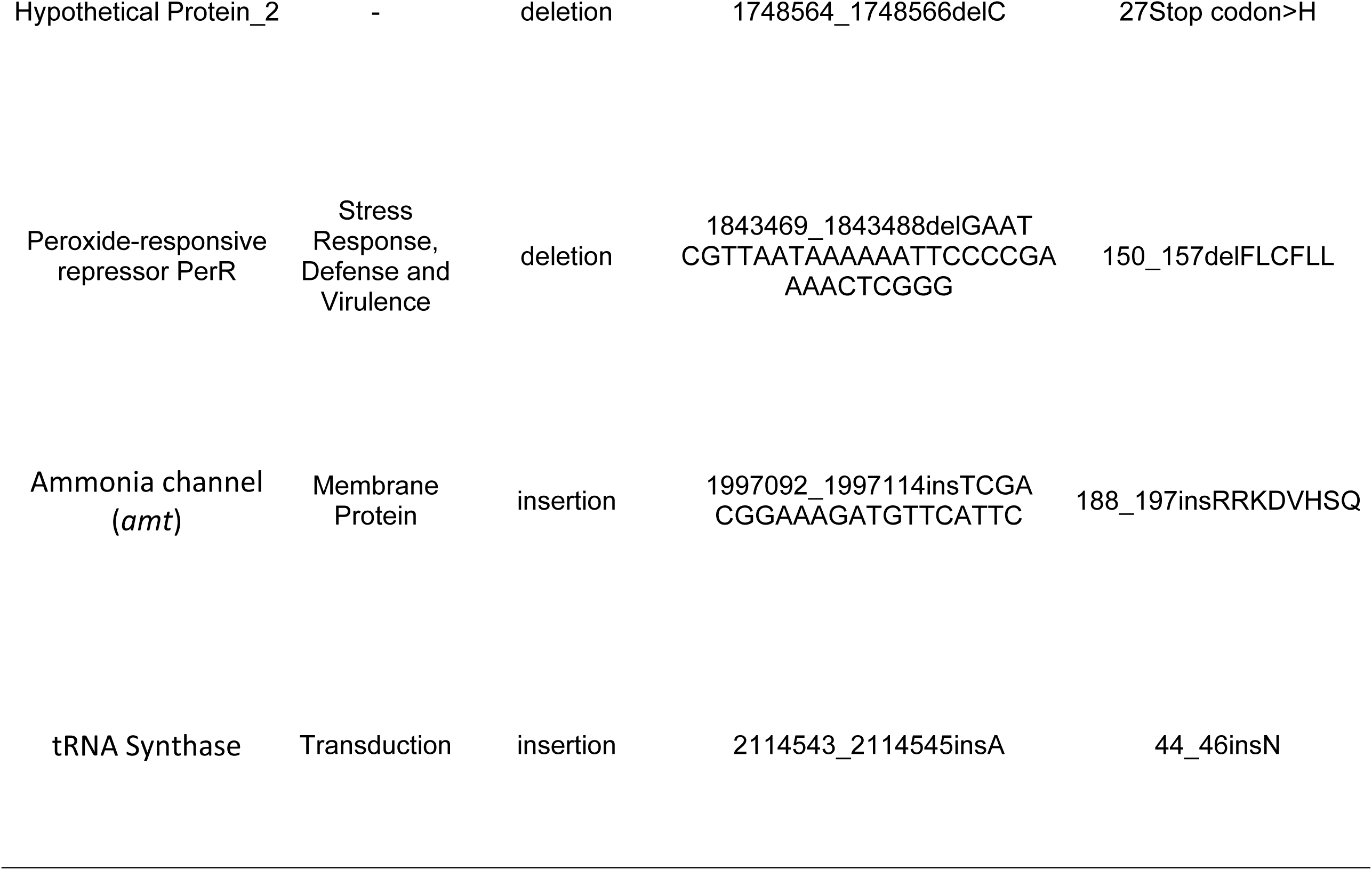
Substitutions of *Lactobacillus reuteri* Rifampicin-resistant mutant Milk-adapted strain.

## DISCUSSION

We showed that an intestinal *L. reuteri* isolate subjected to 150 passes in milk gained proficiency to grow in this substrate but lost aptitude to grow in MRS medium and possibly in murine intestines (table 1). Our results are in agreement with previous studies showing that adaptation to a new ecosystem reduces the fitness in the original ecosystem, a phenomenon known as antagonistic pleiotropy (22, 24, 26). This finding is also in agreement with previous reports indicating that commercial probiotic bacteria (originated from intestines), failed to colonize mammalian intestines (27). We showed that even low passage probiotics may have reduced the ability to colonize intestines.

A total of 13 mutations were observed in bacteria subjected to milk incubation; non-synonymous mutations (potential positive selection) was observed in metabolic genes (putative amino acid permease *yhd*G; sugar phosphatase *ybiV*). The permease YhdG is an exporter for branched-chain amino acids in *B. licheniformis* (28). Mutations of the β′ subunit (*rpo*C) which regulates adaptation to grow in minimal medium (29). Other apparently selected non-synonymous mutations were in the cell wall or cell membrane components (UDP-N-acetylglucosamine-N-acetylmuramyl-(pentapeptide) pyrophosphoryl-undecaprenol N-acetylglucosamine transferase (*mur*G); Putative cell-wall-anchored protein (LPXTG motif). These results are similar to Long Term Serial Passage experiments in *E. coli* in a minimal medium where researchers showed evolutionary convergence in genes that encode proteins with regulatory and metabolic functions (30).

Lactobacilli growing in milk have to endure high concentrations of autogenous organic acids which can be deleterious and bacterial populations subjected to these types of acids can select acid resistant mutants (31). The strain adapted to milk has a non-synonymous mutation in the amino acid permease YhdG gene (branched chain amino acids biosynthesis) which has been showed in an acid tolerant mutant of *Streptococcus mutans* (25).

Peroxide-responsive repressor *per*R gene has a deletion of 34 nucleotides in the strain adapted to milk. PerR is a ferric uptake repressor which regulates genes involved in oxidative stress responses, iron homeostasis and is essential for responses to peroxide (33). A deletion in this repressor gene may cause constitutive expression of these functions in the strain adapted to the milk. This trait may allow bacterial growth in milk, a substrate in which iron is sequestered by lactoferrin and transferrin (34).

A mutation in the *amt* gene (an insertion of 21 nucleotides) was observed in *L. reuteri* adapted to milk, the product of this gene is involved in the ammonium transmembrane transporter activity (35). It has been shown that *L. acidophilus* uses glucosamine-6-phosphate isomerase to process glucosamine 6-phosphate producing fructose 6-phosphate and ammonia and it has been found that *L. acidophilus* grown in milk overexpresses this gene (31). This could be a situation in which the overexpression of the mechanisms that produce ammonium in *Lactobacillus* growing in milk are also selecting mutants with beneficial changes for greater transport of ammonium through the membrane.

Cell-wall-anchored protein (LPXTG motif) show a non-sense substitution that replaced Arginine for a stop codon resulting probably in a loss of its functionality. LPXTG-anchored proteins are covalently attached to the peptidoglycan are some of the outermost structures in *Lactobacillus* and may play an important part in interactions with host components (32) and may no longer be useful when grown in artificial media. Our results are consistent with extensive reductive evolution observed in an *L. bulgaricus* strain (a bacterium originally associated with plants) utilized in yogurt manufacture for a long time (54)

Our failure to recover bacteria from mice intestines may be because rifampin resistance is caused by mutations in the *rpo*B gene; these mutations were observed in our isolates. These mutations cause temperature-sensitive sporulation phenotype (36) and alterations in the expression of global regulons, which control growth, sporulation, and germination in *Bacillus* (37). This subunit is also involved in the binding of the sigma 70 factor (38) to the holoenzyme so that modifications in this subunit could lead to impaired adaptive abilities.

We show that even a few passages in an artificial substrate can select mutants unable to compete with ancestral strains in the original niche. Bacterial antagonistic pleiotropy is a phenomenon that has implications beyond probiotics such as antibiotic resistance and the evolution of environmental opportunistic pathogens.

## MATERIALS AND METHODS

### Isolation and selection of strains of *Lactobacillus reuteri*

The strains used in this study were isolated from the duodenum of a female CD1 mouse (all procedures with mice were previously approved by the Bioethics Committee of the San Francisco University of Quito, Ecuador). A CD1 mouse was euthanized using chloroform, the duodenum was extracted and contents were inoculated onto Man Rogosa Sharpe (MRS) agar and incubated at 37°C for 48 hours under microaerophilic conditions (39). After incubation, they were analyzed morphologically (Gram stain) and biochemically (catalase and oxidase tests) (supplementary table 1). Six *Lactobacillus* spp. isolates were selected and 16S rRNA gene analysis was used to determine bacterial species (40), PCR products were sequenced at Functional Biosciences, Inc., Madison, WI. (supplementary table 1) and sequences were compared to those in GenBank (41).

### Selection of Rifampicin resistant mutants

Rifampicin mutants were selected following the protocols previously described by Miller (42). Four *Lactobacillus reuteri* strains were inoculated into 10 ml of MRS broth at 37°C under microaerophilic conditions for 24 hours, after which bacterial cultures received an additional 10 ml of MRS broth with rifampicin (200μg/ml), for a final concentration of 100μg/ml), incubated for another 24 hours and finally inoculated into MRS agar with rifampicin 100μg/ml) for a final 24-hour incubation. Colonies formed by rifampicin resistant mutants were confirmed with Gram stain and biochemical tests (catalase and oxidase) and antibiotic sensitivity tests (43). Confirmed strains were stored at −80C in BHI medium + glycerol 20%. Fitness loss due to rifampicin resistance was evaluated by culturing together progenitor strains and rifampicin resistant descendants as previously described (44) (supplementary table 2). Finally, we selected a strain LrRR1.2 which showed the least fitness loss to carry out the rest of the experiments (supplementary table 2).

### *Lactobacillus reuteri* Rifampicin-Resistant Strain Milk adaptation

The mutant strain LrRR1.2 was subjected to 150 serial 24h-passes in sterile (autoclaved) cow’s milk incubated at 37°C under microaerobiosis; these number of passes corresponds to approximately 510 generations (45). To rule out potential *Lactobacillus* contamination, PCR amplification and sequencing of *LeuS* gene (46) was carried out at passes: 55, 92 and 120.

### Intestinal colonization capacity assays in murine model

All the procedures performed with the mice were previously approved by the Ethics Committee on the Use of Animals in Research and Teaching at the San Francisco University of Quito. Male and female CD1 mice donated by “Laboratorios Agrocalidad”, Tumbaco, Quito were used. Animals were fed for 7 days with 100 μl of *Lactobacillus reuteri* in culture medium at 2×10^8^ CFU/ml. The animals were separated into three groups: group 1 was fed with *Lactobacillus reuteri* rifampicin-resistant mutant strain (pass 0), group 2 was fed with *Lactobacillus reuteri* rifampicin-resistant mutant strain (pass 150) and the group 3 or control group that was fed only with the culture medium without *Lactobacillus reuteri*. After 15 post-administration days of the treatments, the animals were euthanized as previously described and contents from 10mm of the duodenum were serially diluted and plated in MRS plus rifampicin (47).

### Comparative Growth in MRS broth and Milk

The resulting strains from milk adaptation were taken (ancestor, mutant resistant to rifampicin without milk passes and mutant resistant to rifampicin adapted to the milk) and the individual culture of each one was carried out in both MRS and Milk. Following the methodology of Lenski *et al* (44) the initial population density was quantified and after 24 hours of culture in each medium and relative fitness was defined with the formula W = ln [A_24h_ / A_0h_] / ln [B_24h_ / B_0h_]. Obtaining the growth capacities of each strain in Milk and MRS comparing both the ancestor and the parent before adapting to milk (table 1 and figure 1).

### DNA extraction and Whole Genome Sequencing

*Lactobacillus reuteri* strains sensitive to rifampicin (original ancestor), *Lactobacillus reuteri* rifampicin-resistant mutant strain without milk passes and *Lactobacillus reuteri* rifampicin-resistant mutant strain with 150 milk passes were selected for total DNA extraction using DNAzol™ Reagent, for isolation of genomic DNA from solid and liquid samples (Invitrogen™) following the manufacturer’s protocol (48). The total DNA of the 3 samples was sent to Macrogen Inc., Seoul, the Republic of Korea for genome sequencing, using Illumina Hiseq 2500, 100bp PE.

### Molecular Analysis

The complete genome sequences of the 3 strains were assembled using SPAdes (version 3.13.0). The Raw Data of the sequences were pre-processed before running the hashing. We used paired-ends reads from SPAdes using Mauve version 2.4.0 for the reordering of the contigs (49), based on the complete genome sequence of *Lactobacillus reuteri* DSM 20016 on GenBank (NC_009513.1). The gen annotation of the completed genomic sequences was accomplished in Prokka (version 1.13.3) (50). ProgressiveMauve was used for the alignment of the 3 reordered genomes (51). A Comprehensive Genome Analysis for the characterization of the genome of the three strains, in Patrick version 3.5.22 (52). For the identification of SNPs variants between the milk adapted bacterial genome and its progenitor genome we used the fast bacterial variant calling from NGS reads (Snippy)(53).

## ACKNOWLEDGMENTS

We want to thank the San Francisco University of Quito, especially the College of Biological and Environmental Sciences, COCIBA that funded the research of this study. Also, to the “Laboratorios de Agrocalidad Tumbaco”, Quito that provided the mice for the *in vivo* experiments.

---

**Figure Supp. 1.**
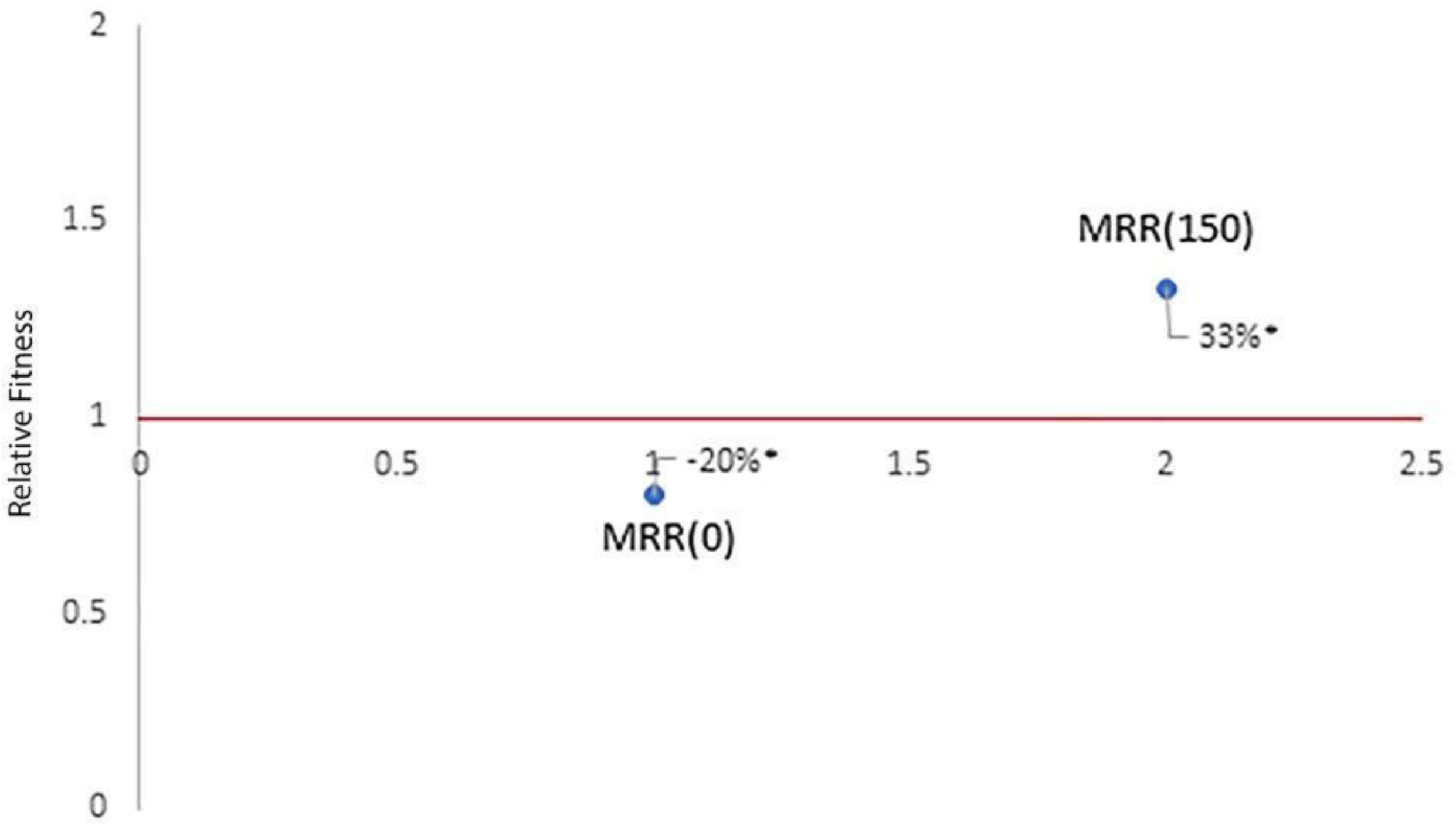
Schematic diagram of the SNPs in *Lactobacillus reuteri* rifampicin mutant resistant strain after 150 passes in milk and *Lactobacillus reuteri* rifampicin resistant mutant prior passes in milk. The colored arrows show the genes where mutations were found.

